# Comparing full variation profile analysis with the conventional consensus method in SARS-CoV-2 phylogeny

**DOI:** 10.1101/2023.08.03.551784

**Authors:** Regina Nóra Fiam, Csabai István, Solymosi Norbert

## Abstract

This study proposes a novel approach to studying SARS-CoV-2 virus mutations through sequencing data comparison. Traditional consensus-based methods, which focus on the most common nucleotide at each position, might overlook or obscure the presence of low-frequency variants. Our method, in contrast, retains all sequenced nucleotides at each position, forming a genomic matrix. Utilizing simulated short reads from genomes with specified mutations, we contrasted our genomic matrix approach with the consensus sequence method. Our matrix methodology accurately reflected the known mutations and true compositions, demonstrating its efficacy in understanding the sample variability and their interconnections. Further tests using real data from GISAID and NCBI-SRA confirmed its reliability and robustness. As we see, the genomic matrix approach offers a more accurate representation of the viral genomic diversity, thereby providing superior insights into virus evolution and epidemiology. Future application recommendations are provided based on our observed results.

## Introduction

In conventional approaches to sequencing the genome of viruses, a consensus sequence is determined^1,2^. The consensus sequence represents the most frequent nucleotide at each position following short-read alignment. There have been attempts to align a large number of consensus sequences with each other, thus obtaining frequencies assignable to each position.^3^ However, one sequence alone does not necessarily accurately represent the composition of the sample, as it represents only a single sequence. Meanwhile, a sample comprises a population of many viruses, with different variants potentially carrying various mutations. This form of reduction results in a loss of information, potentially distorting sample similarity when comparing consensus sequences, and neglecting the “within-host diversity” that characterizes the viral population within a host.

Numerous strategies have been developed to maintain this variety.^4^ Notably, most efforts are documented in the HIV literature, referring predominantly to an approach known as profile sampling. The common factor in these methods is that they assign a frequency to each nucleotide (or codon) following read alignment, creating what constitutes the patient’s HIV profile. The effectiveness of this approach can be enhanced by creating multiple samples from a single frequency table and determining the within-host distribution using these developed profiles through repeated sampling.^5^ The thinking here goes beyond consensus, shifting away from focusing on specific strains. However, we utilize one sample per patient, with our primary interest lying in the relations between these samples, rather than the distribution of strains within a sample.

To address this, we aim to explore how a more accurate depiction of the relationship between individual genomic sequences can be achieved. In this method, the comparison of genome matrices forms the basis for determining the degree of similarity, as opposed to consensus sequences that represent only a single sequence. Each row of the genome matrix represents a genomic position, while the columns indicate the frequency of individual nucleotides. As we have increased the number of dimensions in this approach (since each base now has its own), we expect to lose less information in the comparison process.

We will examine the efficacy of the two methods using simulated sequencing results, then compare the findings and examine which scenario provides a more accurate picture of the known relationships. Finally, we will conduct similarity studies according to the two methods using actual coronavirus sequencing data.

## Materials and Methods

### Bioinformatic analysis of the simulated reads

First, we generated synthetic samples based on the Table 1. In each sample, we precisely defined the mutations that define the variants present in the simulation^6^, as well as the coverage of each variant. For this, we used the `randomreads.sh` tool from BBMap.^7^ We used FASTA files prepared based on the individual variants as reference genomes, generating paired-end reads with a length of `150` base pairs and the desired `coverage` value. We combined the paired-end reads into a single file. The *sequence. f asta* file is the reference genome of SARS-CoV-2, which we indexed with BWA (Version: 0.7.17-r1188), and aligned the generated FASTQ files.^8^ We transformed the resulting .*sam* file into a .*bam* file using the `samtools view-Sb` command.^9^ Then we sorted the .*bam* file using the `samtools sort` command and indexed it using the `samtools index` command, creating the .*bai* files. The consensus sequence was created by using `bcftools consensus`. Firstly, we generated the `mpileup` file using the `bcftools mpileup - Ou -f` tool with the reference genome and the .*bam* file. Afterwards, we utilized the command ‘bcftools call –ploidy 1 -mv -Oz -o’ to generate the ‘vcf’ file. This file was then employed to create the consensus using the command ‘bcftools consensus -f’.^10^

**Table 1.** The table represents the true composition (coverage) of the simulated sample mixtures.

| sample | cheesy | glorious | mushroom | perfume | slinky |
| --- | --- | --- | --- | --- | --- |
| 1 | 50 | 100 | 150 | 200 | 250 |
| 2 | 250 | 50 | 50 | 10 | 10 |
| 3 | 30 | 20 | 100 | 150 | 70 |
| 4 | 10 | 200 | 40 | 30 | 10 |
| 5 | 100 | 100 | 100 | 100 | 100 |
| 6 | 10 | 20 | 15 | 500 | 5 |
| 7 | 40 | 120 | 0 | 10 | 200 |
| 8 | 30 | 500 | 70 | 0 | 20 |

The idea for the genome matrix was given by the Position Probability Matrix used for characterizing sequential motifs.^11^ We used the `Biostrings`, `GenomicAlignments`, `tidyverse` packages in R for the `.bam` files.^12–14^ Essentially, the genome matrix made a .*csv* file from the .*bam* files, where each row contained the counted number of nucleotides aligned at the given positions.

### Sources of the real data

In addition to the simulations, we also carried out the comparison on real data. For this, we had to find samples for which the raw FASTQ file can be found in the NCBI-SRA database^15^ and the consensus in the GISAID database^16^ as shown in Table 2. We selected the reads in such a way that there should be at least 20-fold coverage in at least 0.95 of the genome. Additionally, we worked with paired-end sequencing WGS data. From the SRA, we aligned the reads downloaded with the `SRA Toolkit prefetch` to the reference COVID genome using the `bowtie2` tool.^17^ We use the `bowtie2-build` command to index the reference genome, then we align the reads (paired-end) and thus obtain .*sam* files. The `samtools view -Sb` command converts the .*sam* file into binary .*bam* format, which we similarly sorted and then indexed as in the case of synthetic reads. The genome matrix was also obtained here based on the .*bam* files by counting the nucleotides per position.

**Table 2.** Real data in different databases.

|  | GISAID ID | SRA ID |
| --- | --- | --- |
| 1. | *EPI_ISL_584662* | *ERR4692877* |
| 2. | *EPI_ISL_584663* | *ERR4693014* |
| 3. | *EPI_ISL_584664* | *ERR4693156* |
| 4. | *EPI_ISL_664742* | *ERR4892128* |
| 5. | *EPI_ISL_665153* | *ERR4891061* |
| 6. | *EPI_ISL_665157* | *ERR4890337* |
| 7. | *EPI_ISL_665158* | *ERR4891869* |
| 8. | *EPI_ISL_665166* | *ERR4890532* |
| 9. | *EPI_ISL_665169* | *ERR4891959* |
| 10. | *EPI_ISL_665198* | *ERR4890285* |
| 11. | *EPI_ISL_665199* | *ERR4892877* |
| 12. | *EPI_ISL_665203* | *ERR4891675* |
| 13. | *EPI_ISL_665231* | *ERR4891211* |
| 14. | *EPI_ISL_665232* | *ERR4893062* |
| 15. | *EPI_ISL_665233* | *ERR4892295* |
| 16. | *EPI_ISL_665636* | *ERR4892461* |
| 17. | *EPI_ISL_665848* | *ERR4891212* |

### Comparing the effectiveness of the two methods

The question was to determine whether the relationships or similarities between the samples could be better determined by comparing consensus sequences or by comparing genome matrices. For this, we created distance matrices between the consensus sequences for the 8 samples, similarly for the genome matrices. We first aligned the consensus sequences with each other in R using `AlignSeqs()`, and to visualize this we used `BrowseSeqs()`. Then, we determined the distance matrix for this with the `DECIPHER DistanceMatrix()` function.^18^ We used the default parameters, that is, we calculated with Euclidean distance. We determined the distance matrix from the genome matrices in Python using the `scipy.spatial distance.euclidean()` function.^19^ The *D*_*i, j*_ matrix element contains the distance between the *i*-th and *j*-th genome matrices. We use Euclidean distance as a metric. Accordingly, the distance between two matrices, i.e., an element of the distance matrix, is obtained with the following formula:

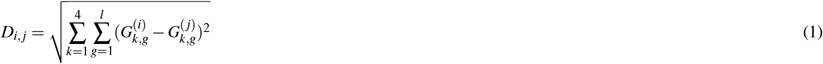

In the above formula, the notations 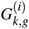 and 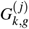 represent the values of the *i*-th and *j*-th genome matrices at the *k*-th base and *g*-th position.

We also created a distance matrix for the true compositions, the result of which also became an 8×8 matrix. This way, we obtained three 8×8 distance matrices. However, before comparing these, we normalized them. We calculated the Frobenius norm^20^ as follows:

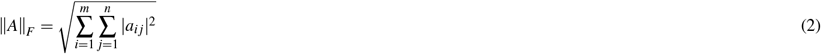

where *A* is a matrix of size *m×n*, and *a*_*i j*_ are the elements of the *i*-th row and *j*-th column of matrix *A*.

### Comparison of the distance matrices

For the comparison of the different methods, we used the distance matrices as a basis. Firstly, we determined the distance (Euclidean) between the distance matrices pairwise, as shown below.

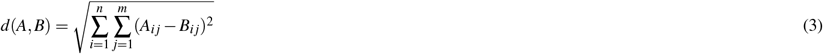

Where *A* and *B* are two *n* × *m* dimensional distance matrices, *A*_*i j*_ and *B*_*i j*_ are elements in the same position.

Secondly, we determined the Pearson correlation between the distance matrices pairwise, which measured the linear relationship between the data, but not necessarily the causal relationship. In Python, we imported the necessary packages using the command `from scipy.stats import pearsonr`, and calculated the degree of correlation between each pair of distance matrices.^19^ However, before that, we ‘flattened’ the matrices for analysis using the `flatten()` function. The `pearsonr` function not only provides the correlation coefficient but also a number called a ‘p-value’, which indicates the probability of whether the correlation between the two samples is real or just a result of chance. In addition, we visualized the relationships expressed by the distance matrices with heat maps and dendrograms. For the heat maps, we used the Python `seaborn` and `matplotlib` packages. We used the `dendrogram` and `linkage` functions from the `scipy.cluster.hierarchy` module to create dendrograms, based on the already prepared distance matrices.

### The role of entropy in the search for polymorphisms

In a large portion of the genome, there is a nucleotide that occurs significantly more frequently at a given position than the others.^21^ In these cases, the frequency of the other nucleotides is typically non-zero, possibly due to statistical or sequencing errors. It’s crucial to note the distortion when normalizing the genome matrix, assigning relative frequencies to each base. Therefore, a variant with high AF at a highly covered position is characterized identically to potential noise at a less covered spot. Hence, there is a need for deep sequencing data to distinguish noise from low AF variants. A valuable adjunctive method involves generating an entropy sequence for the genome matrix, providing the “information” content in different parts of the sequence. The entropy of the genome sequence at certain positions indicates how varied the bases are, i.e., how likely we are to encounter a given base at that position. Calculating entropy helps identify conservative and diverse areas of the genome sequence. Entropy can be calculated using the following formula:

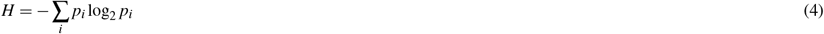

where H is the entropy, and *p*_*i*_ is the probability (frequency) of the i-th base at a certain position. In the present case, the entropy can be calculated as follows:

1. First, calculate the probability of occurrence (normalized frequency) of each base along the rows, i.e., at every position:

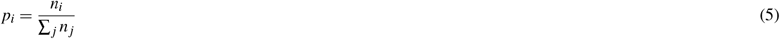

where *n*_*i*_ is the number of occurrences of the i-th base, and summation is carried out over all possible bases (*j*). We already had these values during the normalization of the genome matrix.
2. Calculate the entropy based on normalized frequencies using (4).

## Results and discussion

### Simulation

We denoted *d*(*D*_*GM*_, *D*_*cons*_) as the distance between the distance matrix of the genome matrices and the distance matrix of the consensuses. *d*(*D*_*GM*_, *D*_*true*_) represents the distance between the distance matrix of the genome matrices and the distance matrix of the true compositions per sample. Finally, *d*(*D*_*true*_, *D*_*cons*_) denotes the distance between the distance matrix of the true compositions and the distance matrix of the consensus sequences. Using these notations, the following results were obtained:

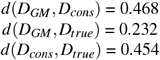

It can be seen that in terms of distances, the reality is better captured by the genome matrix than the consensus-based approach. The results of comparing our matrices gave us the following in terms of Pearson correlation.

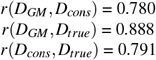

The ‘p-value’ is not specifically mentioned because in every case it was less than 10^*−*15^.

As seen in Figure 1, compared to the true compositions, the consensus approach oversimplifies the results. Seeing the composition of the outlier sample based on Table 1, we can see that in this sample, the variant with the fantasy name *perfume* is dominant. The variant definitions show that there are two deletions among the mutations defining the variant, in addition to the SNPs. Their locations are 11287 (G after deletion of GTCTGGTTTT), and 22298 (A after deletion of AGAAGTTATTTGACTCCTGGTG). Variants that did not contain deletions were *cheesy, glorious*. Hence, it logically follows that sample 6 manifests the most significant deviation from those samples (2,4,7,8) in which these deletion-free variants were predominantly observed. Only in the case of sample 6 did the deleted section make it into the consensus sequences. This is clearly visible at the locations of deletion in Figure 2.

**Figure 1.**
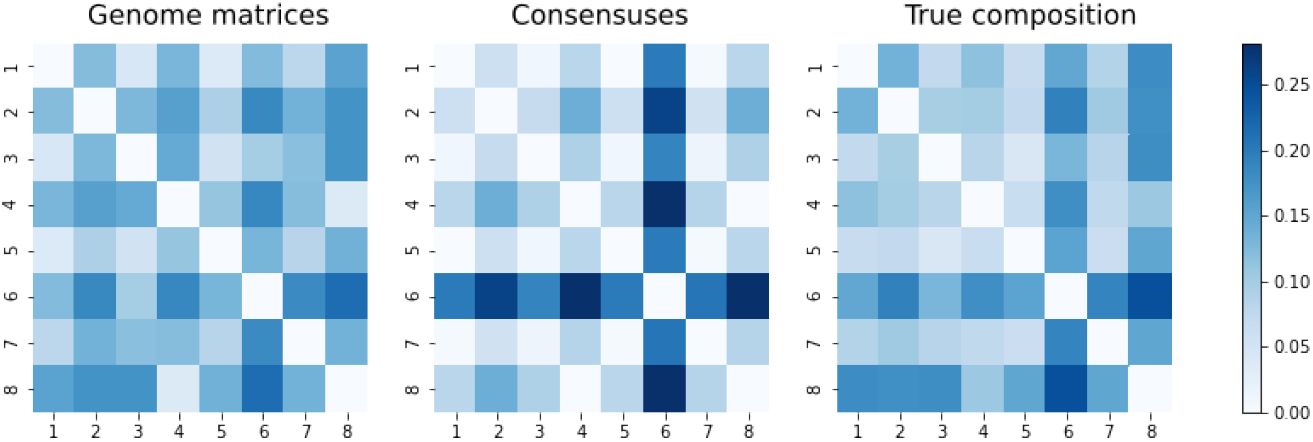
Visualization of the three approaches. The numbers on the axes refer to the order of the simulated sequencing samples listed in Table 1.

**Figure 2.**
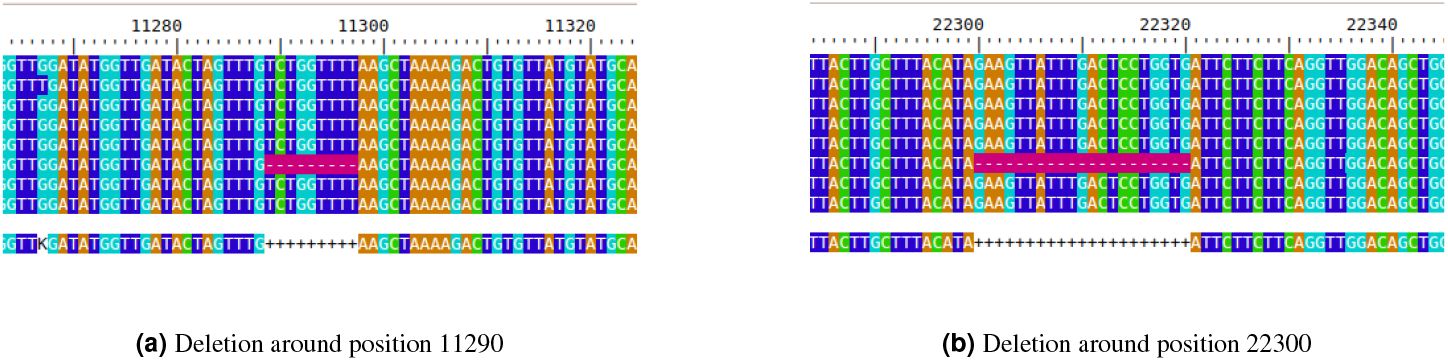
Deletions around the genome marked with ‘-’ in purple

Despite every sample containing a variant that had one of the deletions alongside the SNPs, this only appeared in the consensus sequence of sample 6. The reason for this is that this variant was present with a sufficiently high AF in that sample. In the case of the genome matrix approach, as we could see on the heatmaps, sample 6 was less of an outlier compared to the others. From this, we concluded that the genome matrix is able to take into account deletions with lower AF during comparison.

A dendrogram is a tree-structured diagram that applies hierarchical clustering as seen in Figure 3. In this case, the synthetically created mixed samples are placed on the horizontal axis (x-axis), while the distances can be seen on the vertical axis (y-axis). From Figure 3 we can make the following observations.

**Figure 3.**
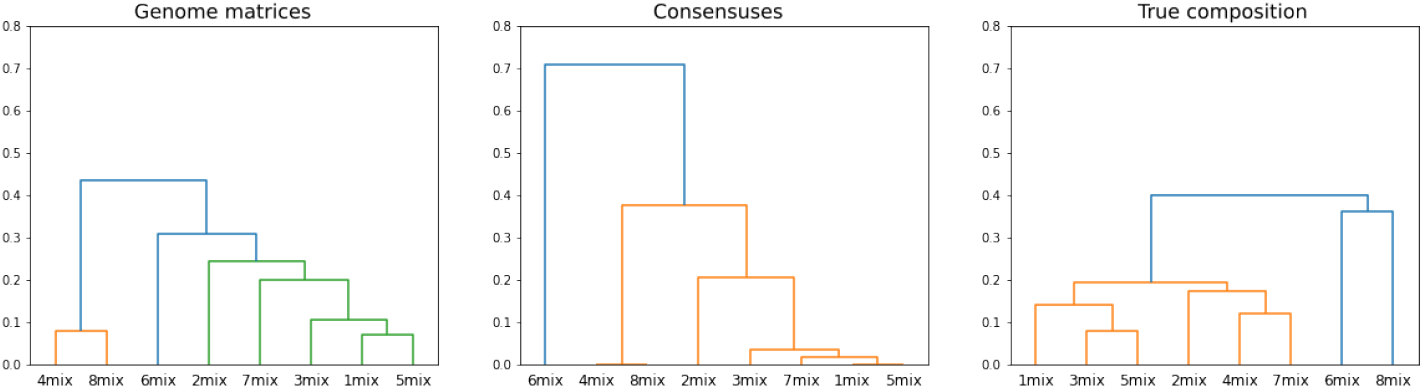
Visualization of the dendrograms of the genome matrix, consensus-based approach, and the true composition’s distance matrices. It can be seen that compared to the true composition, the consensus-based approach amplifies or smoothens the differences, whereas based on genome matrices, we do not obtain such outliers in the relationships.

- According to the dendrogram of the true compositions, two main clusters can be observed. This is understandable if we look at the known composition ratios (Table 1), as in the case of samples 6 and 8, one variant accounts for more than 0.8 parts of the sample.
- According to the true compositions diagram, despite their differences, the samples are at most 0.4 distance from each other. The genome matrix approach slightly amplified this, while the consensus-based procedure nearly doubled it.
- The genome matrix approach performs much better than the consensus-based approach, according to the dendrograms, if a dominant variant contains a deletion segment. In the case of samples 7 and 3, such a variant is dominant in the sample. The consensus-based approach “leveled out” the differences in these samples, resulting in a falsely decreased distance.
- The genome matrix and the consensus-based approach provide similar results in the case where the dominant variants in the samples do not contain a deletion and the other variants are relatively evenly represented. Sample 2 was also like this.
- When a variant containing a deletion segment is dominant, and the other variants (with or without deletions) occur in the sample with a low AF, the consensus method distorts the dendrogram. This is particularly noticeable in the case of sample 6, where the genome matrix does not show such early branching, which is in line with our true composition observations.

### Real data

We downloaded prepared FASTA files from the GISAID database, so it was not necessary to create a consensus sequence from the raw reads as in the case of simulation data. As the sequence lengths in GISAID do not necessarily match, we first performed the MSA on the 17 sequences and the reference genome in R before determining the distance matrix. Both distance matrices were normalized in a similar way for comparability, using Frobenius normalization. Subsequently, we determined the distance and Pearson correlation coefficient of the two distance matrices, similar to the simulations, with the difference that now we did not have the “ground truth” composition distance matrix available.

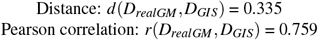

Comparing this with the simulation results, we see that the distance, in this case, is a tenth smaller, but the Pearson correlation between the two matrices is similar. To get a more precise picture of the relation between the two methods, we also compared the two distance matrices using heatmaps (Figure 4).

**Figure 4.**
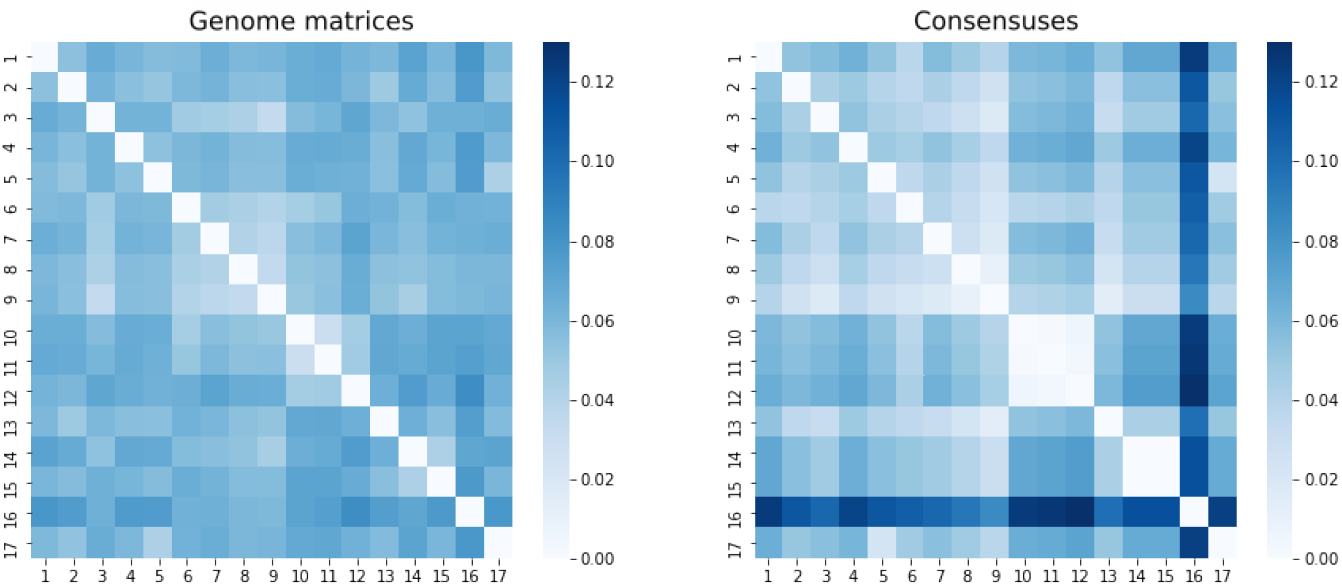
Visualization of the similarity of real data using heatmaps

We saw a similar pattern in the simulation results (Figure 1) as in Figure 4. There was also a tendency that distances were smaller in the case of genome matrices, thus the differences between individual samples. On the other hand, in the consensus-based approach, some samples were very separated from others. According to the lesson of the simulation, this was due to variants containing dominant deletions. In Figure 4, a standout sequence (16th sample: `EPI_ISL_665636, ERR4892461`) can be seen based on the distances from the consensus-based approach.

Figure 5. contains several deletion sections (5), only one of which is present in other samples. Furthermore, the other samples do not contain a deletion section that can only be detected in it.

**Figure 5.**
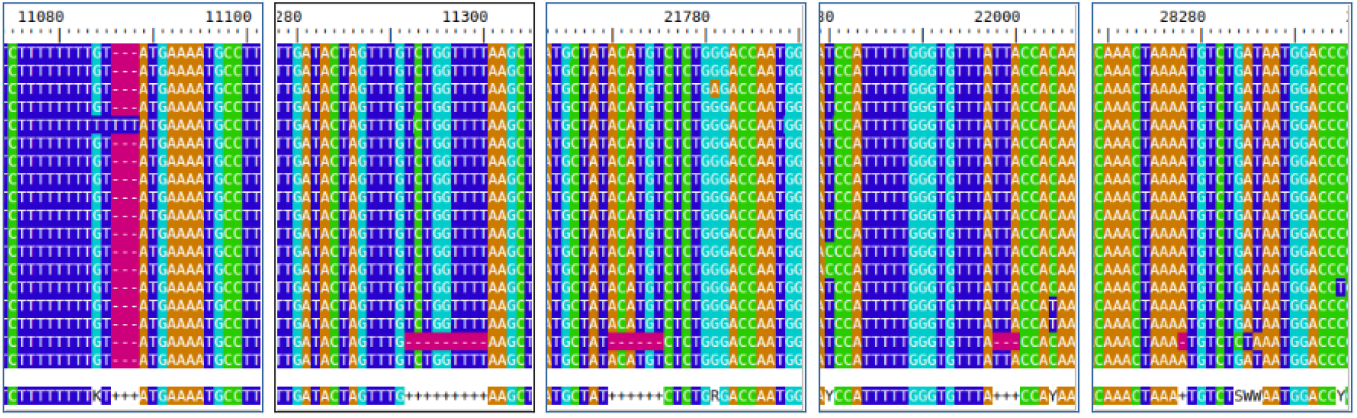
Deletions along the `EPI_ISL_665636` genome marked with ‘-’ in purple

We also looked at these regions based on the genome matrix results. Two trends seemed to be moving on the chart, so we made 2 columns in Figure 6. It is observable that the genome matrix methodology also detects comparable deletion sections, like the consensus-based approach. However, it’s important to note that the frequency of nucleotide hits at these locations doesn’t reduce to a complete zero. This is well visible in the right column as well. In the positions of deletion, an opposite trend appears in this column. In these positions, the number of individual nucleotide hits increases among the lower number of hits. This can also mean that in addition to mis-sequencing, a significant increase can be found in these positions in terms of “secondary” nucleotides. There won’t be much difference between the most common and the second most common nucleotide, as each is relatively low compared to the genomic environment.

**Figure 6.**
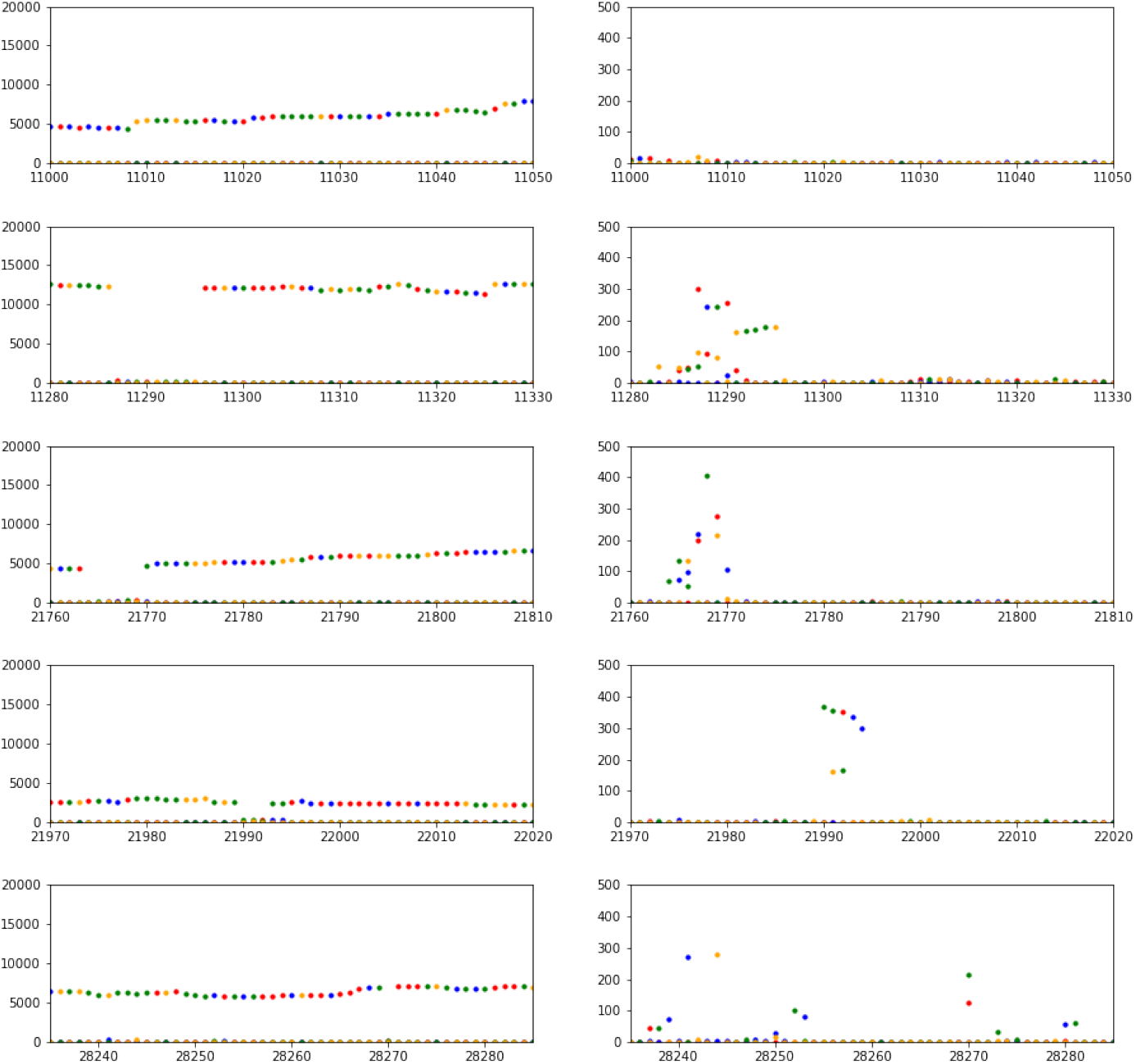
Zooming into the deletion sites within the `ERR4892461` genome. In the left column, it can be seen that “holes” appear in the number of nucleotides at the deletion sites. On the right, zooming into these positions, we can see that there are also alternative nucleotides here, which however, stand out from the noise of the non-deletion sites.

This may mean that in addition to the variant containing a large AF deletion, there are several variants with smaller AFs and these genomic positions are more variable in the sample. However, before we search for these high entropy genomic regions from an information theoretical point of view, we examine the relationship between the samples with dendrograms on Figure 7.

**Figure 7.**
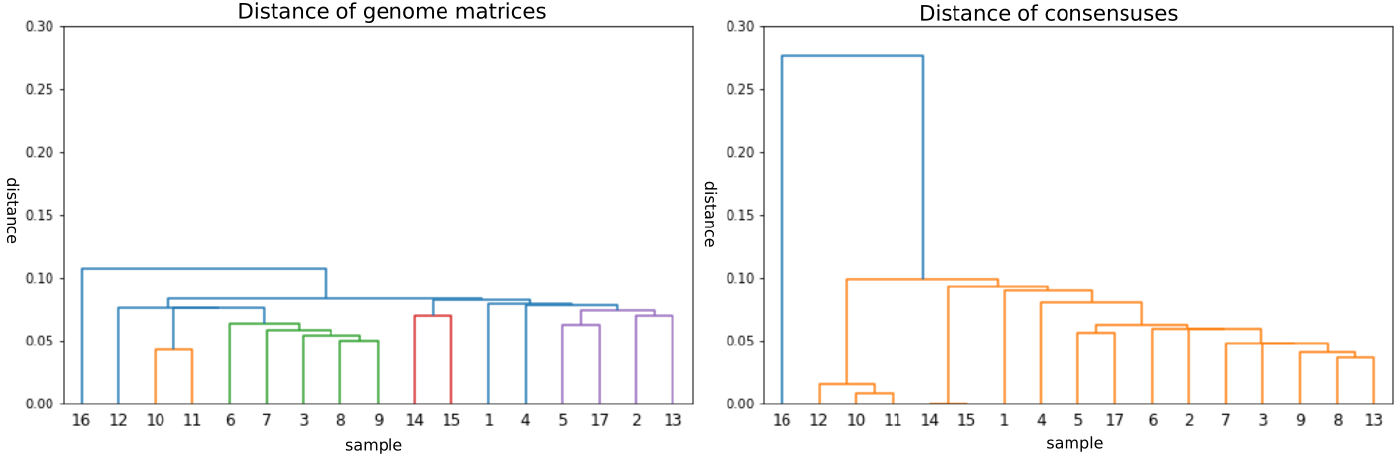
The 16th sample greatly differs from the others, but according to the genome matrix, it shows similarity with several samples. The consensus-based approach organizes the samples into fewer clusters. Accordingly, we concluded here, similar to what was observed in simulations where the true composition was known, that the representation of variability decreases when using consensus.

### Entropy sequences

When detecting deletions, and plotting the total number of nucleotides, we get similar figures as in Figure 6. In such instances, it’s easier to notice low AF variants because they suddenly become dominant as the variant with the highest AF contains some mutation. In the case of normalized genome matrices, these low AF variants also appear, but in a distorted form, since these are relative ratios. Therefore, we used the concept of entropy for a more effective exploration of polymorphisms along the genome, as this method was applicable regardless of alignment. Entropy allows for the measurement of variability and distribution of nucleotides within certain sections, making it easier to identify low AF variants. These could potentially be linked to key features of the examined genome.

For the **real data**, a low entropy value indicates that there is little variability at a given position, and the bases are likely to be the same with a high probability. High entropy values indicate that the bases are diverse and their probabilities are more evenly distributed. Examining entropy values allows for the mapping of the conservative and diverse areas of the sequence. Our data clearly show that for sample 16, around 11300, a higher entropy (higher information content) section is indeed noticeable, which is present in this sample to this extent only. This defining mutation might also be present in others, though this is not clear from the consensus. However, according to the information sequences formed from genome matrices, it is there with low AF. The increased entropy along the genome section in question indicates this.

Figure 8 shows that the entropy of the `ERR4693014` sample is outstandingly high throughout the genome compared to the other genomes. The reason for this could be that it likely contained a mixture of several variants in almost equal measure at the time of sequencing. In this case, even if several strains are present, the consensus sequence will be a single one, making it understandable why it is located so far from the `ERR4892461` sample on dendrograms (Figure 7). In the latter case, the distribution of information content along the genome is less uniform. This could indicate that a dominant strain was present in the sample, whose consensus sequence differed from that of the `ERR4693014` sample, which uniformly incorporated a mixture of many strains.

**Figure 8.**
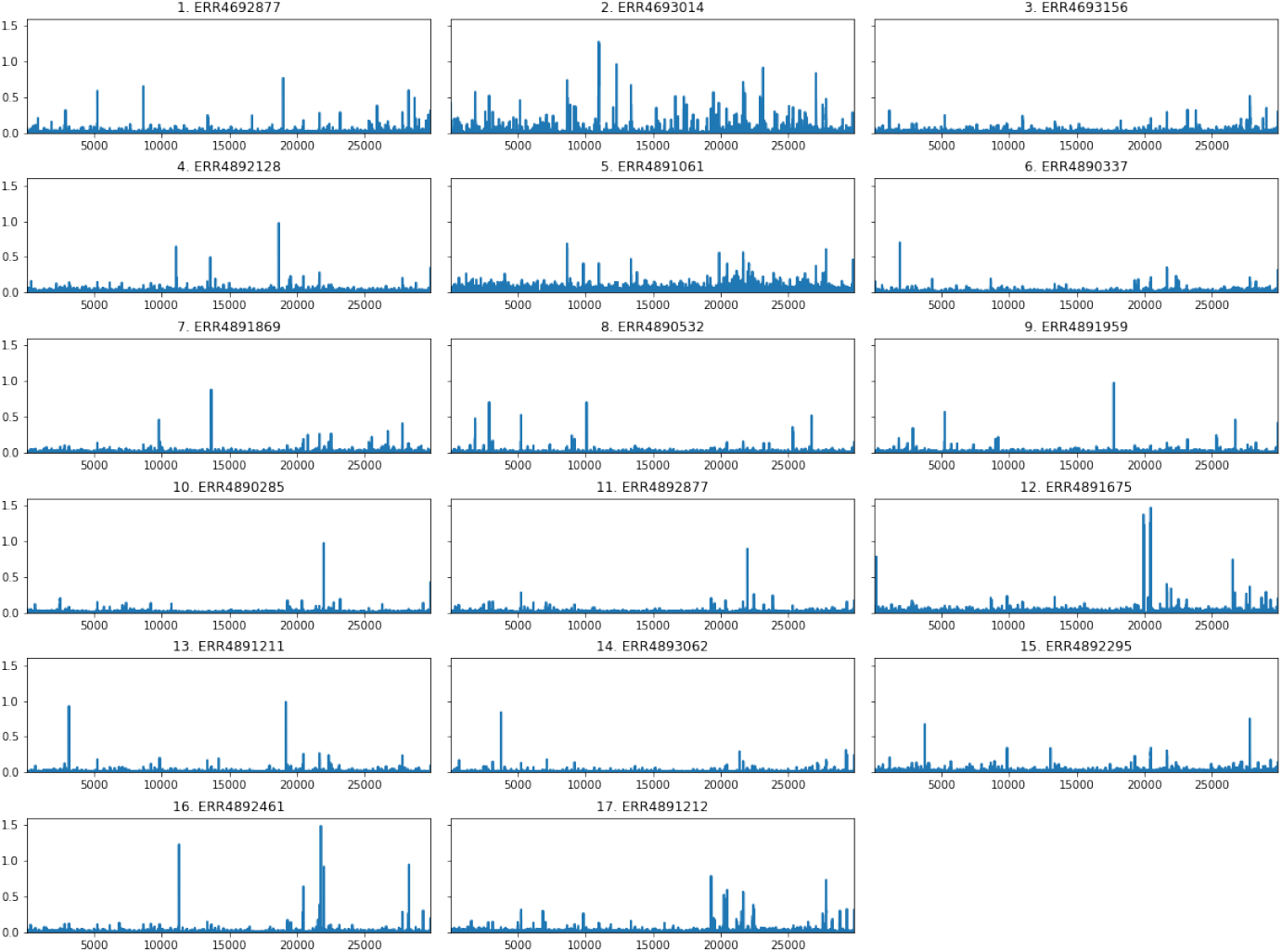
Entropy sequences along the entire genome: visualization of the information content of genomes.

In the **simulation**, we utilized complete knowledge of sample compositions and mutations. We compared consensus sequences and genome matrices, both reliant on Euclidean distance matrices. After Frobenius normalization, we found the genome matrix method superior to the consensus-based approach.

The key takeaways are the following:

- The genome matrix’s distance matrix was more alike the ‘ground truth’, hence more realistic.
- The Pearson correlation between the genome matrix and the true composition distance matrix was higher than that with the consensus, indicating a stronger connection between initial and post-processing data.
- Visualization via heatmaps (Figure 1.) showed that the consensus approach overemphasizes inhomogeneities in mixing ratios, potentially skewing phylogenetic tree generation. No similar distortion was observed in genome matrices.

High entropy denotes high variability, indicating a surprising or high-information content base type. Visualizing entropy values unveils a base entropic background due to sequencing errors or statistical noise. Polymorphisms, with their peaks rising in high entropy, differentiate from this background. The method’s advantage lies in its applicability to various mutation types, be it SNP or deletion/insertion.

This method enables efficient identification of genome anomalies, with the genome matrix approach playing a pivotal role. This allows for more realistic sample comparisons even without precisely determining specific variants and their ratios. It contributes to accurately unveiling phylogenetic and evolutionary correlations in the realm of viruses, thereby increasing the efficiency and reliability of such research.

## Declarations

### Ethics approval and consent to participate

Not applicable.

### Consent for publication

Not applicable.

### Availability of data and material

The datasets used and/or analyzed during the current study are available from the corresponding author upon reasonable request.

### Author contributions statement

All authors read and approved the final manuscript.

## References

1. Mardis, E. R. Next-generation dna sequencing methods. Annu. Rev. Genomics Hum. Genet. 9, 387–402, DOI: https://doi.org/10.1146/annurev.genom.9.081307.164359 (2008).

2. Miller, J. R., Koren, S. & Sutton, G. Assembly algorithms for next-generation sequencing data. Genomics 95, 315–327, DOI: https://doi.org/10.1016/j.ygeno.2010.03.001 (2010).

3. Caraballo-Ortiz, M. A. et al. Tophap: rapid inference of key phylogenetic structures from common haplotypes in large genome collections with limited diversity. Bioinformatics 38, 2719–2726 (2022).

4. Gribskov, M. & Veretnik, S. [13] identification of sequence patterns with profile analysis. In Methods in Enzymology, vol. 266, 198–212 (Elsevier, 1996).

5. Guang, A. et al. Incorporating within-host diversity in phylogenetic analyses for detecting clusters of new hiv diagnoses. Front. Microbiol. 12, 803190 (2022).

6. The UK Health Security Agency. Variant definitions. https://github.com/phe-genomics/variant_definitions/tree/main/variant_yaml. Accessed on 24/05/2023.

7. Bushnell, B. BBMap: a fast, accurate, splice-aware aligner. Tech. Rep., Lawrence Berkeley National Lab.(LBNL), Berkeley, CA (United States) (2014). https://sourceforge.net/projects/bbmap/.

8. Li, H. Aligning sequence reads, clone sequences and assembly contigs with BWA-MEM. arXiv preprint arXiv:1303.3997 DOI: https://doi.org/10.48550/arXiv.1303.3997 (2013).

9. Li, H. et al. The sequence alignment/map format and SAMtools. Bioinformatics 25, 2078–2079, DOI: https://doi.org/10.1093/bioinformatics/btp352 (2009).

10. Danecek, P. et al. Twelve years of SAMtools and BCFtools. Gigascience 10, giab008. DOI: https://doi.org/10.1093/ gigascience/giab008 (2021).

11. Schones, D. E., Sumazin, P. & Zhang, M. Q. Similarity of position frequency matrices for transcription factor binding sites. Bioinformatics 21, 307–313, DOI: https://doi.org/10.1093/bioinformatics/bth480 (2005).

12. Pagès, H., Aboyoun, P., Gentleman, R. & DebRoy, S. Biostrings: Efficient manipulation of biological strings (2022). R package version 2.64.1, https://bioconductor.org/packages/Biostrings,.

13. Lawrence, M. et al. Software for computing and annotating genomic ranges. PLoS Comput. Biol. 9, e1003118. DOI: https://doi.org/10.1371/journal.pcbi.1003118 (2013).

14. Wickham, H. et al. Welcome to the tidyverse. J. Open Source Softw. 4, 1686, DOI: https://doi.org/10.21105/joss.01686 (2019).

15. Leinonen, R., Sugawara, H., Shumway, M. & International Nucleotide Sequence Database Collaboration. The sequence read archive. Nucleic Acids Res. 39, D19–D21, DOI: https://doi.org/10.1093/nar/gkq1019 (2010).

16. Shu, Y. & McCauley, J. GISAID: Global initiative on sharing all influenza data–from vision to reality. Eurosurveillance 22, 30494, DOI: https://doi.org/10.2807/1560-7917.ES.2017.22.13.30494 (2017).

17. Langmead, B. & Salzberg, S. L. Fast gapped-read alignment with Bowtie 2. Nat. methods 9, 357–359, DOI: https://doi.org/10.1038/nmeth.1923 (2012).

18. Wright, E. S. Using DECIPHER v2.0 to Analyze Big Biological Sequence Data in R. The R J. 8, 352–359 (2016).

19. Virtanen. SciPy 1.0: Fundamental Algorithms for Scientific Computing in Python. Nat. Methods 17, 261–272, DOI: 10.1038/s41592-019-0686-2 (2020).

20. Weisstein, E. W. Frobenius norm. https://mathworld.wolfram.com/FrobeniusNorm.html.

21. Shannon entropy. https://www.hiv.lanl.gov/content/sequence/ENTROPY/entropy_readme.html.

